# RNase E endonuclease activity and its inhibition by pseudoridine

**DOI:** 10.1101/2021.05.23.445298

**Authors:** Md. Saiful Islam, Katarzyna J. Bandyra, Yanjie Chao, Jörg Vogel, Ben F. Luisi

## Abstract

The conserved endoribonuclease RNase E dominates the dynamic landscape of RNA metabolism and underpins control mediated by small regulatory RNAs in diverse bacterial species. We explored the enzyme’s hydrolytic mechanism, allosteric activation, and interplay with partner proteins in the multi-component RNA degradosome assembly. RNase E cleaves single-stranded RNA with preference to attack the phosphate located at the 5□ nucleotide preceding uracil, and we corroborate key interactions that select that base. Unexpectedly, RNase E activity is impeded strongly when the recognised uracil is isomerised to 5-ribosyluracil (pseudouridine), from which we infer the detailed geometry of the hydrolytic attack process. Kinetics analyses support models for recognition of secondary structure in substrates by RNase E and for allosteric auto-regulation. The catalytic power of the enzyme is boosted when it is assembled into the multi-enzyme RNA degradosome, most likely as a consequence of substrate channeling. Our results rationalize the origins of substrate preferences of RNase E and illuminate its catalytic mechanism, supporting the roles of allosteric domain closure and cooperation with other components of the RNA degradosome complex.

## INTRODUCTION

RNase E, a key bacterial endoribonuclease of ancient evolutionary origin, has multifaceted activities critical to organism fitness, including the turnover of mRNA, maturation of precursors of tRNA and rRNA, processing and degradation of small regulatory RNAs, and rRNA quality control (Mackie 2013; 1998; Bandyra, Wandzik, and Luisi 2018). Once cleaved by RNase E, an mRNA becomes committed to an irreversible fate of rapid deconstruction; but at the same time, the enzyme can contribute to an orderly genesis of structured RNAs from precursors that circumvents destructive pathways, provided that those species pass quality control checks. The enzymatic activity of RNase E, which appears to be nuanced, serves as a key determinant of cellular RNA lifetime in cells. Its substrate preferences and encounter rate with RNA impact on transcript lifetime *in vivo* and are of interest for elaborating a potential code that could define cellular RNA fate.

Decades of analysis of RNase E activity indicate that there is no simple sequence code for its substrates *per se*, but instead a strong preference to cleave within single-stranded regions enriched in A or U (Chao et al. 2017; Del Campo et al. 2015; Kime et al. 2010; 2014; Mackie 2013). Global RNA target analyses performed both *in vivo* and *in vitro* identify uracil positioned to the 3□ side adjacent to the nucleotide of the scissile phosphate (the +2 position) as a strong signature for RNase E activity (Chao et al. 2017). For many substrates in both destructive and maturation pathways, the enzyme is activated by transformation of the 5□ end of the substrate from a triphosphate normally found on nascent transcripts, to a monophosphate found on processed species (Mackie 2013). For other substrates, the status of the 5□ end is not so critical for RNase E action (Baker and Mackie 2003; Clarke et al. 2014; Kime et al. 2014), and for these ‘5□ end bypass’ substrates, other features such as secondary structure of the RNA appear to be important. Secondary structure also contributes to recognition of sites for cleavage in both degradative and processing pathways (Richards and Belasco 2021; Bandyra, Wandzik, and Luisi 2018; Updegrove et al. 2019).

The critical endonuclease activity of RNase E is encompassed within the highly conserved amino-terminal domain (NTD) (Figure 1a), which corresponds to roughly half the mass of the protein. Structural studies of this domain have provided insight into the origins of substrate recognition and 5□-end dependent activation (Callaghan et al. 2005; Bandyra, Wandzik, and Luisi 2018; Koslover et al. 2008) (Figure 1a). Key structural motifs include an RNA binding S1 domain and a 5□-sensor that can read the chemical status of the RNA 5□-end (Figure 1a). The recognition of the 5□-end triggers a conformational switch that maneuvers the S1 domain to clamp onto substrates and present them in the active site with geometry favorable for hydrolytic attack. A zinc-coordination motif links covalently protomers into a dimer that is the fundamental catalytic unit, and a dimer-of-dimer results through self-complementary association of a small domain that is evolutionarily related to KH RNA binding (Pereira and Lupas 2018) (Figure 1a). A vestigial RNase H-like subdomain carries no known catalytic function but has been observed to cooperate with the KH-like small domain to recognize duplex structures in substrates and help present adjacent single-stranded regions to the proximal active site (Bandyra, Wandzik, and Luisi 2018) (Figure 1a). Surprisingly, single atom substitutions in a conserved pocket of this domain drive the enzyme into a hyperactive state (D26N, D28N, D338N) (Bandyra, Wandzik, and Luisi 2018; Updegrove et al. 2019). These observations support a model in which the RNase H-like domain auto-regulates the activity of the enzyme by influencing the energetics of domain closure (Bandyra, Wandzik, and Luisi 2018).

**Figure 1.**
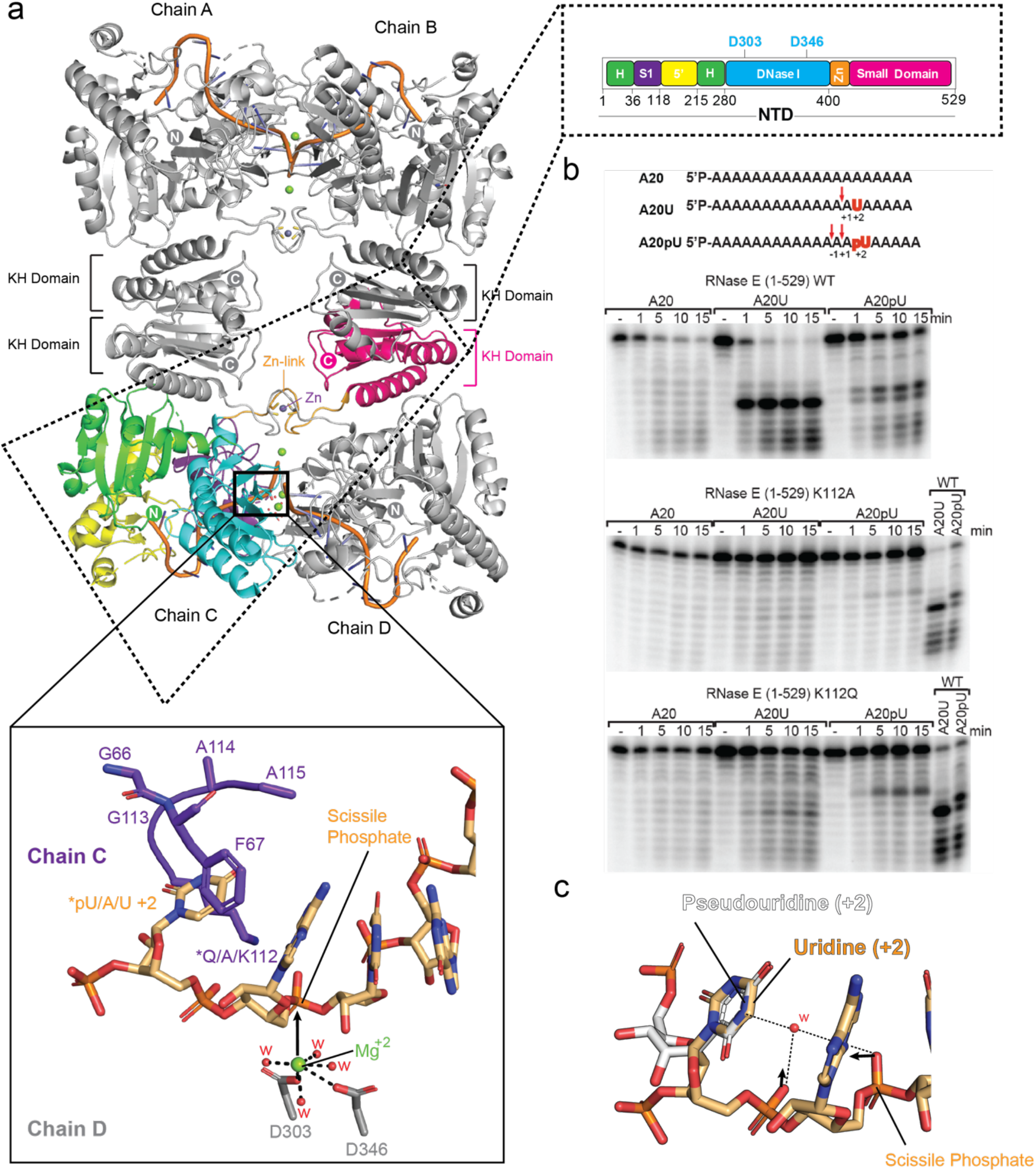
Role of K112 and uracil +2 in substrate interaction. (a) RNase E NTD forms a tetramer, and the cartoon represents the crystal structure of protomers in complex with RNA (PDB: 2C0B) (Callaghan et al. 2005) showing RNA binding domains (as colored in domain architecture), active-site residues Asp^303^ and Asp^346^, active-site Mg^+2^ (green spheres) coordination of which is fulfilled by water molecules (red spheres), Zn-link comprising two cysteine residues Cys^404^ and Cys^407^ (orange and gray sticks) and one Zn^+2^ atom (purple sphere). Inset shows the organisation of RNA substrate binding on the S1 domain of RNase E and the scissile phosphate in the RNA (light orange) bound in the active site on the interface of two protomers presented for the hydrolytic attack by the waters associated with magnesium ion (Mg^++^, green sphere); the Mg^++^ is coordinated by D346 and D303; the U+2 is bound in the uracil pocket formed by amino acids K112, G113, A114 and A115. (b) Cleavage assays of RNase E. Cleavage of 20-mer poly A (A20), poly A with U at position 15 (A20U) or poly A with pseudouridine (Ψ) at position 15 (A20Ψ) by wild-type RNase E NTD (top panel), K112A (middle panel) and K112Q (bottom panel). The substrate was 5’ end-labelled and the products resolved on by denaturing urea-PAGE. The timepoints of the reactions are annotated above the gels.(c) Representation of hydrogen bond mediated hydration organisation at site of pseudouridine. pU/A/U/ +2 = pseudouridine/adenine/uracil at +2 position of the scissile phosphate.

The carboxy-terminal half of the protein, which is predicted to be intrinsically disordered (Aït-Bara, Carpousis, and Quentin 2015; Aït-Bara and Carpousis 2015; Callaghan et al. 2005), provides the scaffold to assemble protein partners into the RNA degradosome complex (Bruce et al. 2018; Bandyra et al. 2013; Bandyra et al. 2018). Through the cooperation of its components and recruitment of RNA chaperones such as Hfq, the RNA degradosome is the central machinery in *Escherichia coli* and many other species for processing of structured precursors and turnover of RNA. RNase E also contains a short amphipathic α-helical domain that interacts with the *E. coli* inner membrane, and the resulting membrane localisation of the degradosome adds a spatial layer to post-transcriptional gene regulation (Hadjeras et al. 2019). Two RNA binding sites in the RNase E c-terminal domain, referred to as AR1 and AR2, cooperate with RhlB to assist in substrate unwinding and remodelling (Khemici and Carpousis 2004; Garrey et al. 2009; Leroy et al. 2002; Chandran et al. 2007). The two RNA binding sites, together with RhlB can interact with ribosomes (Tsai et al. 2012) and may enable degradosome to cleave mRNA in support of a proposed scavenging process (Deana and Belasco 2005; Dreyfus 2009). A plausible scenario is that the close proximity of the RNA degradosome to the translational machine prevents the translation of aberrant transcripts and rescues stalled ribosomal assemblies as part of bacterial RNA surveillance.

Open questions remain regarding details of the RNase E catalytic mechanism, and its capacity to act on modified RNA. The interplay with the components of the degradosome on the quantitative activity of the catalytic domain also have not been evaluated. In this report, we explore the activity of RNase E on substrates with pseudouridine, finding that surprisingly the enzyme is very sensitive to this modification. We measured the ribonuclease activity of RNase E and variants that effect substrate recognition, and we explored how the RNA degradosome assembly cooperates with this activity. Taken together, our results provide mechanistic insights into RNase E catalytic mechanism, allostery, and cooperation within the RNA degradosome complex.

## RESULTS

### K112 plays an important role in substrate preference and cleavage by RNase E

Modelling using the crystal structures of RNase E catalytic domain predicts that S1 domain residues K112 and F67 interact with the base at position +2 to orient the single stranded region of the RNA substrate into a favourable geometry at the active site for nucleophilic attack by water (Chao et al. 2017) (Figure 1a). Uracil at the +2 position is favoured by a hydrogen bonding interaction between the amino group of K112 and the exocyclic carbonyl groups that contributes to the sequence preference at that position. The +2 base is also predicted to be sandwiched between the aromatic ring of F67 and the acyl component of the K112 side chain (Figure 1a). It seems likely that the amino group of the K112 may stabilise the charge on the phosphate in the hydrolytic intermediate.

To test the importance of K112, we compared the activities of the purified wild-type and mutant catalytic domain of RNase E NTD using a model single-stranded RNA substrate composed of 20 nucleotide polyA with a single uracil at position 15 (Figure 1b). The time course for the cleavage is shown in Figure 1b, with products resolved on a denaturing gel. At the enzyme:substrate ratios used in these assay conditions, corresponding to multiple turnover conditions, RNase E cleaves efficiently at the phosphate 2-nt upstream of uridine, consistent with the +2U ruler-and-cut mechanism (Chao et al. 2017). The cleavage activity of the uracil-containing substrate is greater compared with the polyA substrate that does not contain uracil (Figure 1b, top panel, compare A20 and A20U). When K112 is substituted with alanine, the enzyme activity and specificity are greatly diminished for the uracil-containing substrate, with more starting substrate remaining over the time course and the degradation pattern resembling a uniform ladder, as distinct from being enriched for a particular species (Figure 1b, middle panel, compare A20 and A20U with top panel). Even the comparatively conservative substitution of K112 with the long polar side chain of glutamine has diminished cleavage preference for the U+2 position (Figure 1b, bottom panel). Based on the crystal structure, the glutamine is predicted to be too short to interact with either the uracil or the phosphate backbone. These results corroborate the importance of the K112 interaction for catalysis and suggest that the hydrogen bonding interaction with either the uracil base, the scissile phosphate, or both are required for optimal activity.

### Pseudouridine impedes RNase E activity and shifts the cleavage site

The substitution of the uracil at position +2 with pseudouridine (Ψ) involves an isomeric transformation of the base and is not expected to impact on the presentation of the hydrogen bonding O2 and O4 of uracil (Figure 1c). However, pseudouridine has a profound effect on the cleavage activity of RNase E (Figure 1b top panel, compare A20U with A20Ψ). Most of the Ψ-containing substrate resists cleavage by RNase E in the course of the experiment. The cleavage that does occur is partially shifted to the -1 position. These findings indicate that the recognition of uracil is not simply due to a hydrogen bonding interaction with the principal substituents of the base, but also depends on the detailed interactions that influence the phosphodiester geometry. Most likely these details affect the hydration pattern of the substrate and the energy required to achieve the conformation that enables development of the enzymatic transition state.

Substitution of K112 with Q changes the cutting pattern for the Ψ containing substrate. In general, substitution of lysine by the polar glutamine is expected to retain capacity for hydrogen bond formation. Despite the reduced sensitivity to RNase E, the preferred cleavage site was moved to the +2 or +3 position relative to the site cut by wild-type enzymes (Figure 1b, middle and bottom panels). The cutting pattern and shift effect are substantially weakened for the K112A substitution, suggesting that activation of hydrolysis requires a long polar side chain at position 112 (Figure 1a). The K112Q substitution perhaps cause the substrate to align differently in the active site so that it is displaced by one or two nucleotides in the 3□ direction compared to the corresponding wild-type complex.

### RNase E catalytic power can be boosted by substitutions at DNase I and RNase H-like domains

Earlier studies showed that the catalytic activity of RNase E is boosted by mutations of conserved, non-catalytic residues in the RNase H-like domain (D26N and D28N) and DNase I domain (D338N) (Figure 2b) (Bandyra, Wandzik, and Luisi 2018). The substitutions are at a distance from the active site but involve areas where the conformation changes associated with apo to substrate-bound states and are likely to impact on allosteric switching of the enzyme (Bandyra, Wandzik, and Luisi 2018) (Figure 2b). We compared the catalytic activit1y of the wild type and hyperactive triple mutant D26N, D28N, D338N (NTD-3M). For substrates, we used the sRNA GlmZ, which is inactivated by RNase E cleavage, and 9S RNA which is a precursor of ribosomal 5S RNA. The enzyme cleaves the 9S mainly at three sites to form the p5S precursor ribosomal RNA product (Figure 2a) (Cormack and Mackie 1992; Christiansen 1988). The 5□ phosphorylation state of 9S RNA can impact on the cleavage events, with the second event having the activating 5□P group present and anticipated to be intrinsically accelerated if the group is read by the enzyme.

**Figure 2.**
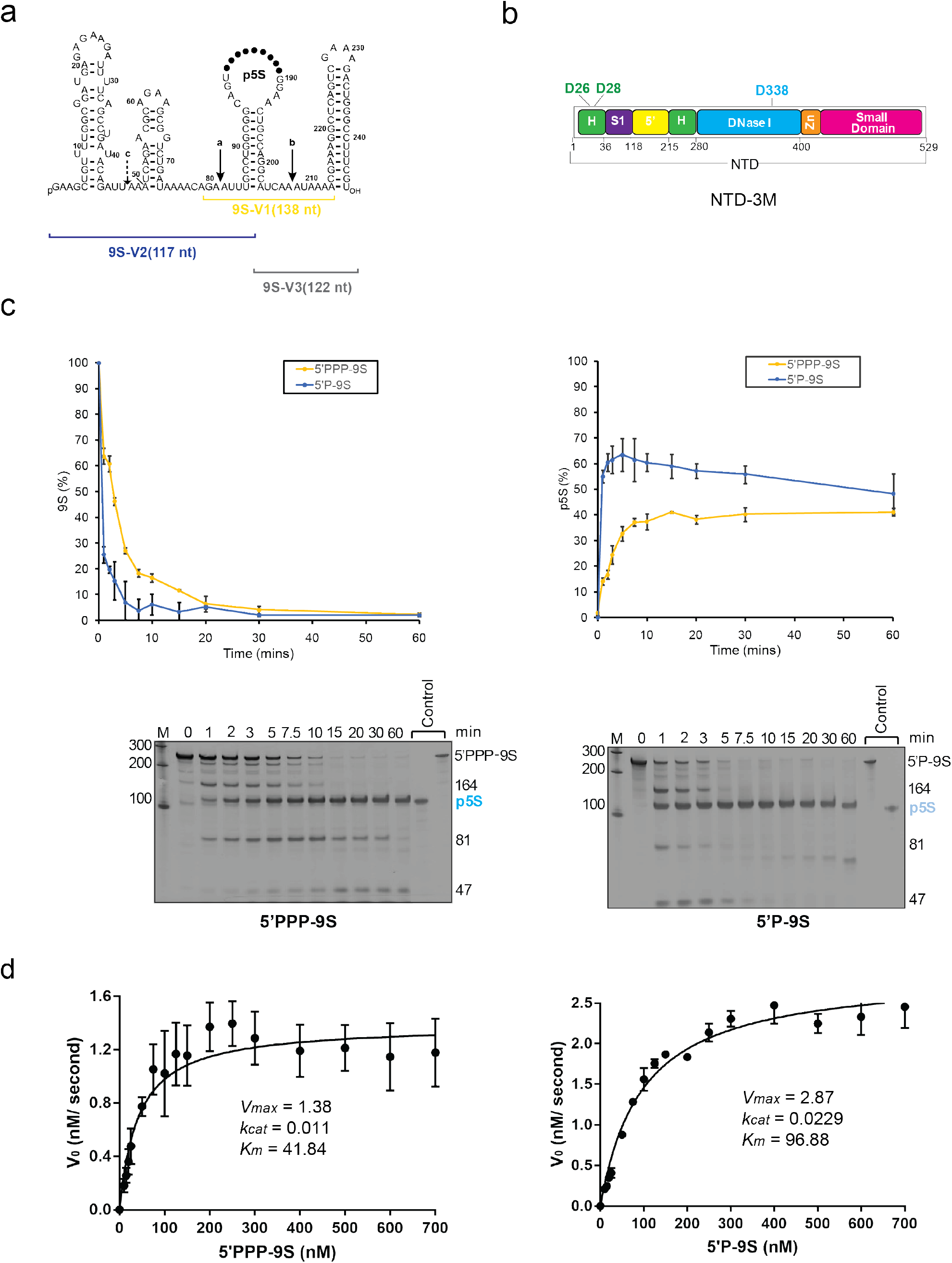
Mutations in the RNase H-like domain improve catalytic efficiency of RNase E. (a) 9S RNA schematic showing the predicted structural elements and the “a”, “b”, and “c” cleavage sites for RNase E (Lorenz et al. 2011; Christiansen 1988). (b) Domain architecture of RNase E NTD showing the position of three mutations (D26, D28, and D338) that were inserted on RNase H- and DNase I-like domains. (c) Plots showing time course assay of 9S (5’-monophosphorylated and 5’-triphosphorylated) cleavage tested against RNase E NTD with triple mutations (D26N, D28N, D338N) (NTD-3M); insets are the urea-denaturing gels showing cleavage products of 9S. (d) Michaelis-Menten plots used to determine the kinetics parameters of 9S cleavage catalyzed by NTD-3M. Values represent mean ± standard deviation (n = 3).

Compared to the wild type, NTD-3M showed higher activity for 5□PPP-9S and a boost in catalytic power due to both increased catalytic rate and decreased *K*_*m*_ (Figure 2c and 2d, Table 1). The results support the proposed role of allosteric autoregulation of enzyme activity, in which domain closure helps to pre-organize the active site so that the apparent affinity of the Michealis-Menten complex increases probably by decreasing the energy barrier to capture and engulf the substrate.

**Table 1.** Kinetic parameters for RNA cleavage catalyzed by RNase E catalytic domain and the degradosome assembly

| Enzyme | Concentration (nM) | RNA substrate | Substrate Degradation /Product Formation (%) | $V_{max}(\text{nM.s}^{-1})$ | $K_m(\text{nM})$ | $k_{cat}(\text{s}^{-1})$ | $k_{cat}/K_m(\text{s}^{-1}.\text{nM}^{-1})$ |
| --- | --- | --- | --- | --- | --- | --- | --- |
| NTD | 125 | 5' PPP-9S | 95.5/28.6 | 0.90 | 75.44 | 0.0072 | $9.5 \times 10^{-5}$ |
| | | 5' P-9S | 95.6/36.3 | 1.44 | 85.79 | 0.0115 | $13.4 \times 10^{-5}$ |
| | | 5' PPP- | 97.5/ND | 4.81 | 521.40 | 0.0384 | $7.2 \times 10^{-5}$ |
| NTD-3M | 125 | 5' PPP-9S | 97.3/41.1 | 1.38 | 41.84 | 0.0110 | $26.5 \times 10^{-5}$ |
| | | 5' P-9S | 98.0/48.2 | 2.87 | 96.88 | 0.0229 | $23.5 \times 10^{-5}$ |
| | | 5' PPP- | 99.4/ND | 11.68 | 729.00 | 0.0934 | $12.7 \times 10^{-5}$ |
| Truncated-degradosome | 50 | 5' PPP-9S | 99.6/NA | 1.05 | 61.13 | 0.0210 | $34.3 \times 10^{-5}$ |
| | | 5' P-9S | 100 | 2.26 | 116.30 | 0.0453 | $38.9 \times 10^{-5}$ |
| | | 5' PPP- | 100 | 7.02 | 545.30 | 0.1404 | $25.6 \times 10^{-5}$ |
| Full-degradosome | 25 | 5' PPP-9S | 99.6/51.9 | 2.1 | 95.26 | 0.0839 | $88.0 \times 10^{-5}$ |
| | | 5' P-9S | 100 | 2.32 | 121.10 | 0.0931 | $83.7 \times 10^{-5}$ |
| | | 5' PPP- | 100 | 7.15 | 679.20 | 0.2863 | $42.1 \times 10^{-5}$ |

5' PPP = 5'-triphosphate, 5' P=5'-monophosphate

### Metals in the catalytic mechanism: RNase E active site may recruit one metal in the apo form

The active site, which is present within a DNase I sub-domain of the NTD, bears two conserved aspartate residues (D303 and D346) that recruit magnesium ion to activate a water molecule for nucleophilic attack on the scissile phospho-diester bond (Thompson, Zong, and Mackie 2015; Callaghan et al. 2005) (Figure 1). The binding interactions between RNase E and metal cofactors was evaluated by isothermal calorimetry (ITC) using a variant of RNase E with residue D346 replaced with a cysteine residue, which was reported previously to be catalytically active in presence of Mn^++^, not Mg^++^ (Figure 3a) (Thompson, Zong, and Mackie 2015). Testing the activity of the D346C NTD on two different RNAs, 9S and the small RNA RprA, confirms that the enzyme is active for cleavage only in the presence of Mn^++^ (Figure 3b and 3c). Using isothermal titration calorimetry (ITC) and titrating the mutant enzyme against Mn^++^ yields a *K*_*D*_ for metal binding in the absence of RNA at 17 µM, with associated ΔH = -19.45 kcal/mol and ΔS = -35.4 cal/mol/deg (Figure 3d). The binding profile indicates that one metal ion can be bound by each subunit of the catalytic domain in the absence of substrate.

**Figure 3.**
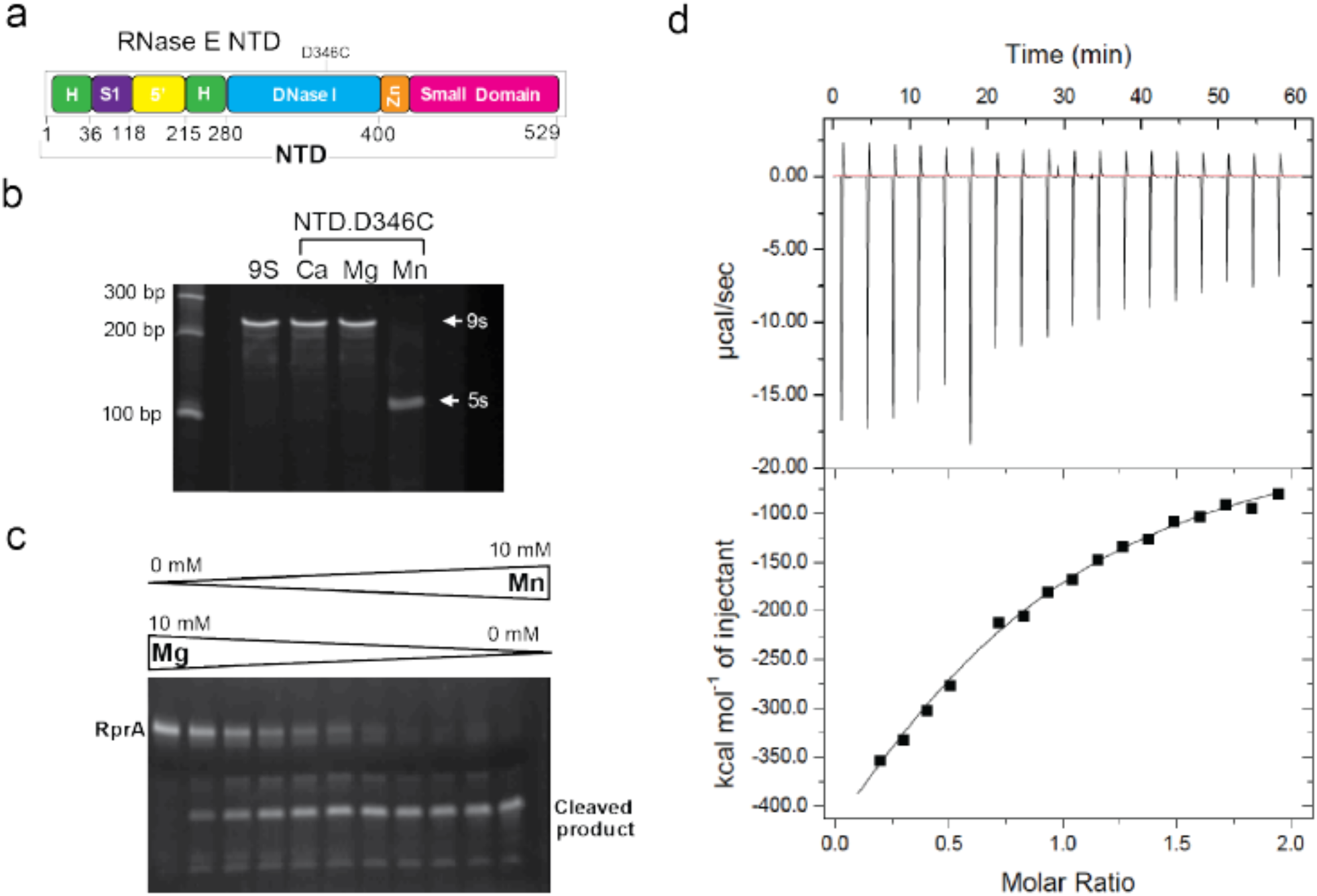
RNase E metal interactions. (a) RNase E NTD variant RneNTD.D346C is catalytically active in presence of Mn^++^ but not any other metal as was seen for processing of 9S and RprA (b,c, respectively). (d) A titration curve of RneNTD.D346C binding to Mn^++^. The *K*_*D*_ is 17 µM for Mn^++^. The titration curve is representative of three experiments.

### Probing RNase E mechanism with unnatural amino acids

To explore interactions between NTD substrates we used derivatives of the protein with the photocrosslinkable amino acid para-azido-phenylalanine (p-AzidoPhe) incorporated at specific positions in the 5□-sensing pocket and duplex-RNA binding site using the amber-supressed system (Chatterjee et al. 2014) (Figure 4a). The substitutions were made singly at residues Met^130^, Ile^139^, Arg^142^, and Tyr^269^ (Figure 4b). Surprisingly, time-course activity assays indicate that formation of the p5S species from 9S is impeded by the substitutions in the 5□-sensing pocket, suggesting that the substitutions may perturb the pocket (Figure 4c). On the other hand, the substitution at the duplex binding surface had little impact on activity (Figure 4c). Exposing NTD p-AzidoPhe derivatives to light at 254 nm in the presence of 9S subdomains did not yield appreciable photocrosslinking directly to the RNA, as detected by mobility shifts in denaturing protein gels (Figure 4d). However, the protein migrated differently in the denaturing gel when uv-illuminated in the absence of RNA, and this may arise from intra-molecular crosslinks (Figure 4d). These results support earlier findings that the NTD undergoes conformational change upon RNA binding (Koslover et al. 2008; Bandyra, Wandzik, and Luisi 2018; Callaghan et al. 2005).

**Figure 4.**
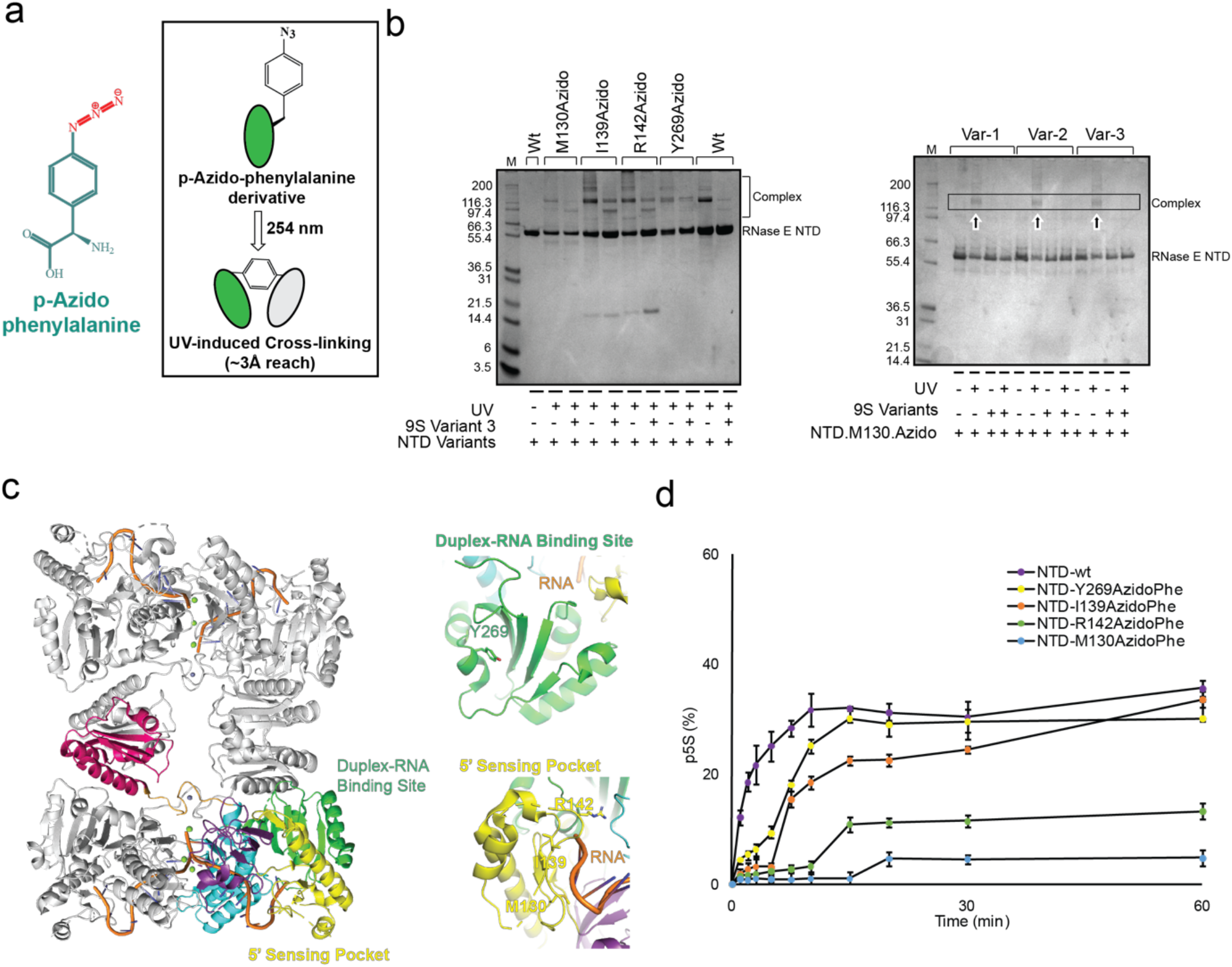
Incorporation of azido-phe into RNase E. (a) Chemical formula of para-azido-phenylalanine (p-AzidoPhe); inset shows p-AzidoPhe to undergo photo-crosslinking to nearby basic residues upon UV exposure at 254 nm. (b) Denaturing protein gels showing p-AzidoPhe derivatives of RNase E NTD form cross-linked product(s) which get interrupted due to the presence of 9S RNA. The p-AzidoPhe did not photo-crosslink to RNA but might have made intra-domain interaction(s) which are lost in the presence of 9S RNA, suggesting a conformational change upon RNA binding.(c) Models of RNase E NTD tetramer with bound RNA at active site, 5’ sensor and the duplex recognition region with insets showing the residues (M130, I139, R142, and Y269) mutated to p-AzidoPhe. The crystal structure of NTD.RNA complex (PDB: 2C0B) was used in this model (Callaghan et al. 2005). (d) Time course assay of p5S production from 9S RNA, processed by p-AzidoPhe derivatives of NTD. Values represent mean ± standard deviation (n = 3).

### Activities of the degradosome for multi-step processing of complex substrates

To explore how RNase E activity is impacted by the degradosome organization, we explored the activity of the isolated catalytic domain with the degradosome for several RNA substrates (Figure 5a). The activity of purified degradosome and the isolated NTD for 9S RNA cleavage was measured using as substrate 9S RNA with 5□-triphosphate (5□PPP-9S) or 5□-monophosphate (5□P-9S). The reaction was first explored with the NTD, which gave the expected p5S product (Figure 5b) and the apparent kinetic parameters for formation of p5S are summarized in figure 5d and Table 1. These are comparable to values reported earlier, and differences are likely due to reaction conditions (Redko et al. 2003; Thompson, Zong, and Mackie 2015; Hadjeras et al. 2019). Comparing the kinetics parameters, the assays revealed significant acceleration to cleave 5□P-9S versus 5□PPP-9S (Figure 5b-5d), corroborating earlier findings that 5□-sensing can contribute to the first cleavage event in 9S processing by RNase E (Mackie 2013; Cormack and Mackie 1992).

**Figure 5.**
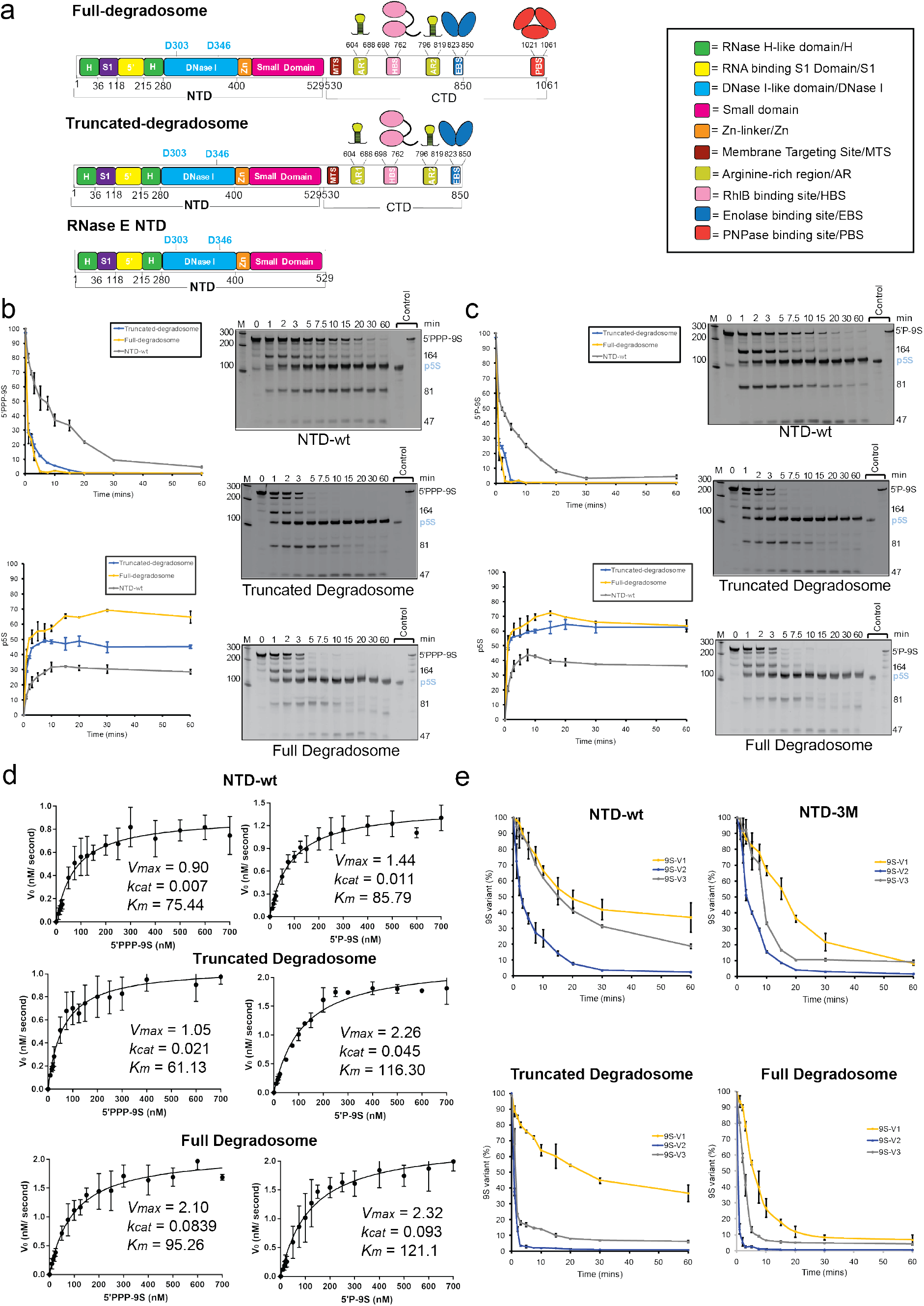
9S cleavage catalyzed by RNase E NTD and degradosome assemblies. (a) Domain architectures of full-degradosome (RNase E, enolase, RhlB, and PNPase), truncated degradosome (RNase E, enolase, and RhlB), and RNase E NTD. (b,c) Time course cleavage assay showing disappearance of 9S RNA and production of the precursor RNA p5S for 9S RNA with (b) 5□-triphosphate (PPP-9S) and (c) 5□-monophosphate (P-9S). (d) Michaelis-Menten curves showing 9Scleavage rate (both 5’-mono- and -triphosphorylation) catalyzed by RNase E NTD, truncated degradosome, and full-degradosome. (e) Time-course cleavage of 9S variants catalyzed by degradosome assemblies. Values represent mean ± standard deviation (SD) (n = 3).

Next, we explored the activity of RNase E within the context of the RNA degradosome. We prepared purified recombinant degradosome (comprising RNase E 1-1061, RhlB, enolase, and PNPase) as well as a subassembly comprising RNase E 1-850, RhlB, and enolase (truncated degradosome, TD; Figure 5a). The processing of 5□PPP-9S showed relatively faster cleavage of 9S for TD and full degradosome assemblies compared to the isolated catalytic domain under the same experimental conditions (Figure 5b-5d) and greater catalytic power (*kcat/K*_*m*_), mostly through changes to the apparent *K*_*m*_ (Figure 5d, Table 1). The cleavage rates were also seen to be greater using 5□P-9S as substrate (Figure 5c, 5d). These observations suggest that the degradosome assembly facilitates RNase E activity, most likely through substrate capture that decrease the effective *K*_*m*_.

### Domain decomposition of 9S and impact on RNase E and degradosome activities

RNase E activity can be directed by structural elements, and for the 9S substrate stem-loop II has previously been shown as the minimal structural requirement needed for RNase E to cleave at the ‘*a*’ site (Figure 2a) (Mackie 2013; Cormack and Mackie 1992). To evaluate the impact of structural elements on the microscopic rate constants for the individual cleavage sites of 9S RNA, we prepared subdomains of 9S with various cut-sites: version 1 has cut-sites ‘*a*’ and ‘*b*’, version 2 has cut-site ‘*a*’, and version 3 has cut-site ‘*b*’ (Figure 2a). Under non-catalytic conditions RNase E NTD can form complexes with the isolated 9S subdomains, presumed to mimic Michaelis-Mention states, as shown by electrophoretic mobility shift assay (results not shown). Cleavage assays show production of the p5S from version 1 and generation of RNA species from versions 2 and 3 for NTD, degradosome and truncated degradosome assemblies (Figure 5e). Compared to 9S, version 1 as substrate gave rise to the comparatively slower appearance of p5S (Figure 5e), perhaps due to the absence of the 5□stem-loop structures (both I and II, Figure 2a). Although it lacks the 3□ terminator hairpin, version 2 seemed to be as good a substrate as 9S RNA, with cleavage by RNase E releasing the 81-base 5□ spacer domain (Figure 5e). An intermediate cleavage rate of version 3 compared to version 2 suggests 5□ stem-loops, stem-loop II in particular, support efficient 9S cleavage (Figure 5e). These results confirm the impact of 9S RNA secondary structures on recognition by RNase E during the processing of 9S RNA.

### Activity of the degradosome in single-step cleavage of the riboregulatory GlmZ

The regulatory RNA GlmZ contributes to homeostasis of glucosamine-6 phosphate (G6P), a precursor in the peptidoglycan synthesis pathway and other metabolic routes (Urban and Vogel 2008; Kalamorz et al. 2007; Göpel et al. 2013). As part of the control network, GlmZ is inactivated by RNase E with the help of the adaptor protein RapZ (Gonzalez et al. 2017; Durica-Mitic and Görke 2019; Kalamorz et al. 2007; Urban and Vogel 2008; Göpel et al. 2013). We explored the effects of RapZ on RNase E mediated cleavage of GlmZ (Figure 6a). Analysis of the cleavage reaction revealed production of the GlmZ-Pro, the cleavage product of GlmZ lacking the *glmS* recognition site (Figure 6b). While GlmZ-Pro was formed in the presence of RapZ for the 5□-PPP-GlmZ form, the 5□P-GlmZ form was rapidly chased into a smaller fragment. In the presence of another chaperone protein Hfq, less GlmZ was processed by the NTD, but the RapZ-specific cleavage product GlmZ-Pro was still formed (Figure 6b, c). The hyperactive version of NTD (NTD-3M) was also found to efficiently generate GlmZ-Pro in the presence of RapZ (Figure 6d).

**Figure 6.**
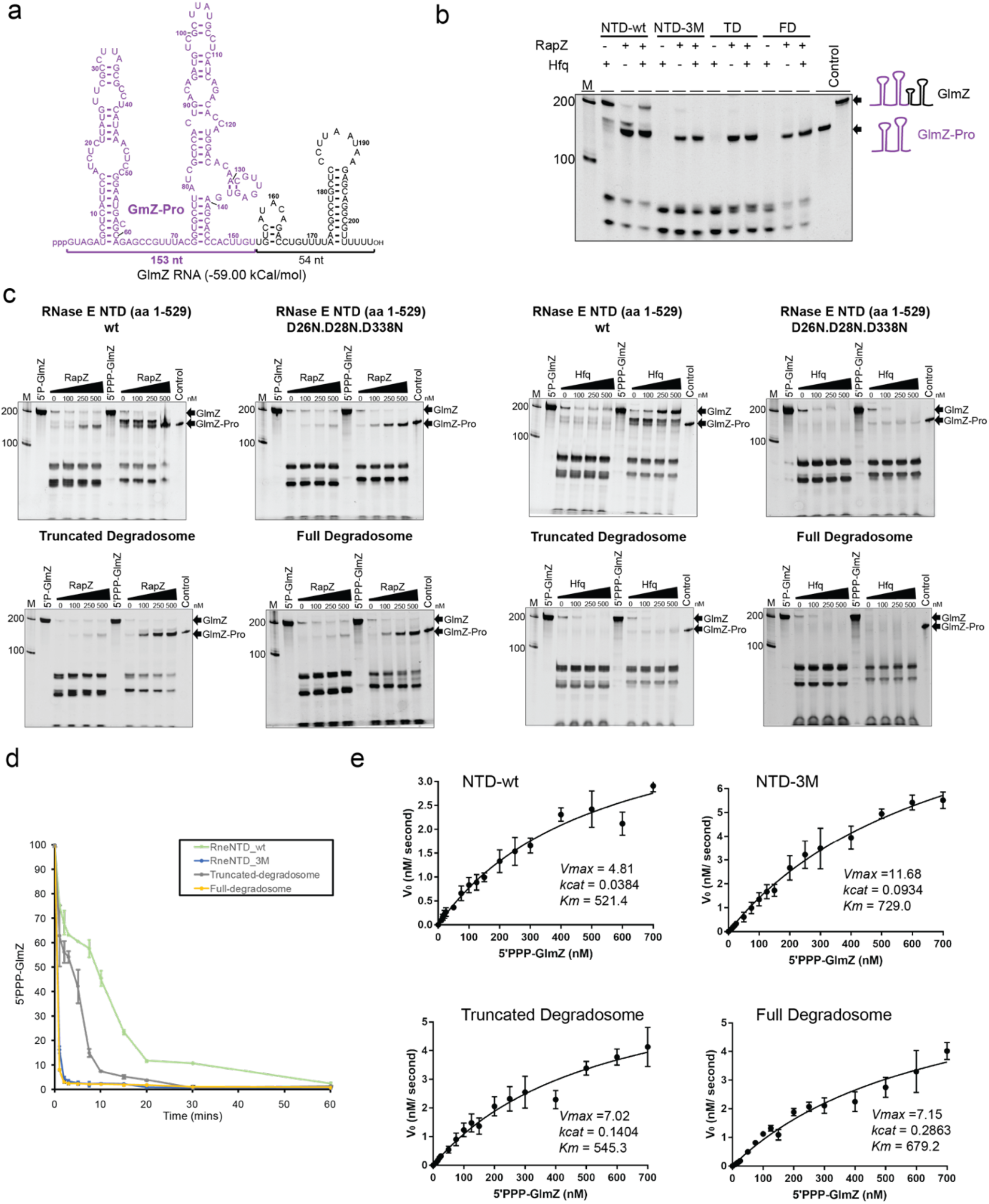
Cleavage of the regulatory RNA GlmZ catalyzed by the RNase E catalytic domain and RNA degradosome assemblies. (a) Secondary structure prediction of GlmZ RNA made by the ViennaRNA Package 2.0 (Lorenz et al. 2011). (b) Single point cleavage assay of GlmZ in presence of RapZ or Hfq or both; please note that the processed form of GlmZ lacking the *glmS* recognition site (GlmZ-Pro) was produced only in presence of RapZ. (c) Urea-denaturing gels showing the cleavage of GlmZ RNA catalyzed by RNA degradosome subassemblies or catalytic domain in the presence of various concentrations of RapZ or Hfq. (d) Time-course quantification of cleavage of GlmZ. (e) Michaelis-Menten plots used to determine the kinetics parameters of GlmZ cleavage catalyzed by RNA degradosome assemblies. Values represent mean ± standard deviation (n = 3).

We determined the kinetics parameters for GlmZ cleavage using degradosome assemblies, revealing faster cleavage of GlmZ by the degradosome and truncated degradosome compared to NTD (Figure 6d and 6e, Table 1). Similar to the findings with 9S RNA (Figure 5), these results indicate that the RNase E mediated cleavage of RNAs can be facilitated in the context of the degradosome assembly.

## Discussion

The half-lives of most *E. coli* transcripts are defined by the activity of RNase E, and sequence and structural preferences for substrates have been identified from *in vitro* and *in vivo* experiments (Clarke et al. 2014; Kime et al. 2014; Chao et al. 2017; Mackie 2013). We explored the catalytic rates of RNase E cleavage of different RNAs to gain further insight into substrate recognition. The cleavage assays with 9S and its truncated versions suggest that RNase E action depends mostly on the secondary structures upstream to the cleavage site, and partly on the secondary structure downstream to the recognition site. Cleavage of all investigated RNAs is influenced by the RNA degradosome assembly. Corroborating earlier findings, mutations in the RNase H-like subdomain boosts hydrolytic activity(Bandyra, Wandzik, and Luisi 2018). A higher reaction rate for the NTD-3M mutant with lower *K*_*m*_^*app*^ and higher *k*_*cat*_ suggest that the RNase H-like and DNase I domains help not only to cleave RNAs but also to release the products quicker than wild type.

The results presented here corroborate the importance of uracil at position +2 with regard to the cleavage site as a key feature of a preferred cleavage site by RNase E and the role of residue K112 in recognising the +2 uracil. Unexpectedly, cleavage by RNase E is strongly impeded when the +2 uracil is substituted with pseudouridine, which is surprising given that this substitution presents only one new hydrogen bonding group on the pyrimidine. The isomerisation of uracil to pseudouridine presents the N1 as a hydrogen bond donor and most likely affects the hydration pattern that will include the phosphate backbone. In most RNA structures, N1 is predicted to interact with the phosphate backbone of both the pseudouridine and the 5□ residue (Charette and Gray 2000). In the context of RNase E catalytic site, this interaction could restrict the backbone conformation at position +2 and disfavour the geometry necessary for catalysis.

Pseudouridine is a commonly occurring modification of tRNA and rRNA in all domains of life (Charette and Gray 2000). The modification of tRNA fragments with pseudouridine has been implicated in translation control in early stages of mammalian embryogenesis (Guzzi et al. 2018). In *E. coli* and other bacteria, the precursors of tRNAs and rRNAs are matured by RNase E cleavage, and the enzyme contributes to quality control of rRNA (Sulthana, Basturea, and Deutscher 2016). As part of the mechanism of quality control, RNase E could hypothetically sense whether the precursors have been properly modified with pseudouridine and destroy those that have not undergone the isomerisation. However, our tests of RNase E activity on tRNAs isolated from cells that are deficient in pseudouridine synthase show that these species, as well as the wild type controls, are resistant to digestion (data not shown). Recent studies suggest that pseudouridine is also prevalent in mRNAs and noncoding RNAs, and that pseudouridylation is regulated by environmental stresses and nutrient availability (Carlile et al. 2014). Differential sensitivity of Ψ to ribonucleases may provide a new mechanism to control RNA stability and/or turnover. Lastly, the results presented here may offer a method to map pseudouridine positions in a sample of RNA through differential sequencing. For example, comparison of RNA sequencing of sample digested with wild type and mutant RNase E (K112Q or K112A) might reveal attenuation of signal for substrates with uridine at +2, but a shift of signal to the -2 or -3 position in the presence of pseudouridine. This could help to pinpoint pseudouridine positions in denatured samples of cell extracted RNA.

The degradosome scaffolding domain of RNase E is predicted to be natively unstructured, and this property has been highly sustained in evolution (Marcaida et al. 2006; Aït-Bara and Carpousis 2015). Recent findings indicate that the disorder property may enable the degradosome to form microscopic condensates in the presence of RNA (Al-Husini et al. 2020; 2018), a property shared with many other RNA binding proteins from all domains of life(Lin et al. 2015; Boeynaems et al. 2018). Enzymatic activities can be concentrated within these bodies, and the environment can affect substrate RNA secondary structures (Guzikowski, Chen, and Zid 2019; Nott et al. 2015). The RNA degradosome from the aquatic Gram-negative bacterium *Caulobacter crescentus* coalesces into nano-scale condensates upon RNA-binding, and these are reversed by RNA turnover (Al-Husini et al. 2020; 2018). Similarly, the membrane associated *E. coli* RNA degradosome forms transient clusters over the membrane during RNA turnover (Moffitt et al. 2016; Strahl et al. 2015). The ribonucleoprotein bodies stimulate RNA decay of target RNAs and complete mRNA turnover, so preventing accumulation of potentially harmful degradation intermediates.

The results presented here show that the catalytic power of RNase E is boosted when the enzyme is assembled into the multi-enzyme RNA degradosome assembly, most likely through substrate channeling. Our observations suggest that the degradosome facilitates RNase E activity, most likely through substrate capture and allostery-mediated acceleration of catalytic rates. We anticipate that the clustering of degradosomes in bodies with liquid-like phase separation further concentrates the enzymatic activities of the machinery and changes the physicochemical conditions that impact on activity. Our results rationalize the origins of substrate preferences of RNase E and illuminate its catalytic mechanism, supporting the roles of allosteric domain closure and cooperation with other components of the RNA degradosome complex.

## Materials and methods

### RNase E NTD expression and purification

RNase E (1-529) wild type and mutants were prepared as previously described (Callaghan et al. 2005; Bandyra, Wandzik, and Luisi 2018). In brief, *Escherichia coli* strain BL21(DE3) was transformed with vector pET16 expressing RNase E (1-529) with an *N*-terminal his_6_-tag. Cultures were grown in 2xTY media supplemented with 100 µg/mL carbenicillin at 37°C, in an orbital shaker set at 220 rpm. The culture was induced between 0.5 to 0.6 OD_600nm_ by adding 1 mM isopropyl-β-thiogalactopyranoside (IPTG) and harvested after 3 hours of incubation by centrifugation at 4200 *g* and 4° C for 30 minutes. Cell pellets were stored as suspension in nickel-column buffer A (20 mM Tris pH 7.9, 500 mM NaCl, 5 mM imidazole, 1 mM MgCl_2_) at -80° C. Once thawed, the cell culture suspension was supplemented with DNase I and EDTA-free protease inhibitor cocktail tablet (Roche) and cells were lysed by passing through an EmulsiFlex-05 cell disruptor (Avestin) for 2-3 times at 10-15 kbar pressure. The lysate was clarified by centrifugation at 35000 *g* for 30 minutes at 4° C and the supernatant was loaded onto a pre-equilibrated HiTrap Chelating HP column charged with nickel ions (GE Healthcare). The column was washed extensively with wash buffer (20 mM Tris pH 7.9, 500 mM NaCl, 100 mM imidazole, 1 mM MgCl_2_), followed by gradient elution of RNase E with elution buffer (20 mM Tris pH 7.9, 500 mM NaCl, 500 mM imidazole, 1 mM MgCl_2_). Fractions containing RNase E were pooled and loaded on a butyl sepharose HP column (GE Healthcare) which previously was equilibrated in high-salt buffer (50 mM Tris pH 7.5, 50 mM NaCl, 25 mM KCl, 1 M (NH)_2_SO_4_). A gradient of a low-salt buffer (50 mM Tris pH 7.5, 50 mM NaCl, 25 mM KCl, 5% glycerol) was used to elute protein. Fractions containing RNase E were pooled, concentrated and loaded onto a size-exclusion column (Superdex™ 200 Increase 10/300, GE Healthcare) equilibrated previously in storage buffer (20 mM HEPES pH 7.5, 500 mM NaCl, 10 mM MgCl_2_, 0.5 mM TCEP, 0.5 mM EDTA, 5% glycerol). The optimal fractions were flash frozen in liquid nitrogen and stored at -80 °C until further use.

### RNase E NTD azido-phenylalanine incorporation and purification

An amber suppressor codon (AUG) was inserted at defined positions of NTD by site-directed mutagenesis. The sequences of the primers used to insert AUG codons are provided in Table 2. Para-azido-phenylalanine (p-AzidoPhe) was inserted in RNase E NTD by co-expressing pET16 carrying *rne* gene corresponding to NTD and pDULE2 carrying *gene* corresponding to orthogonal tRNA synthetase in *Escherichia coli* BL21(DE3) (Chatterjee et al. 2014).

**Table 2.**
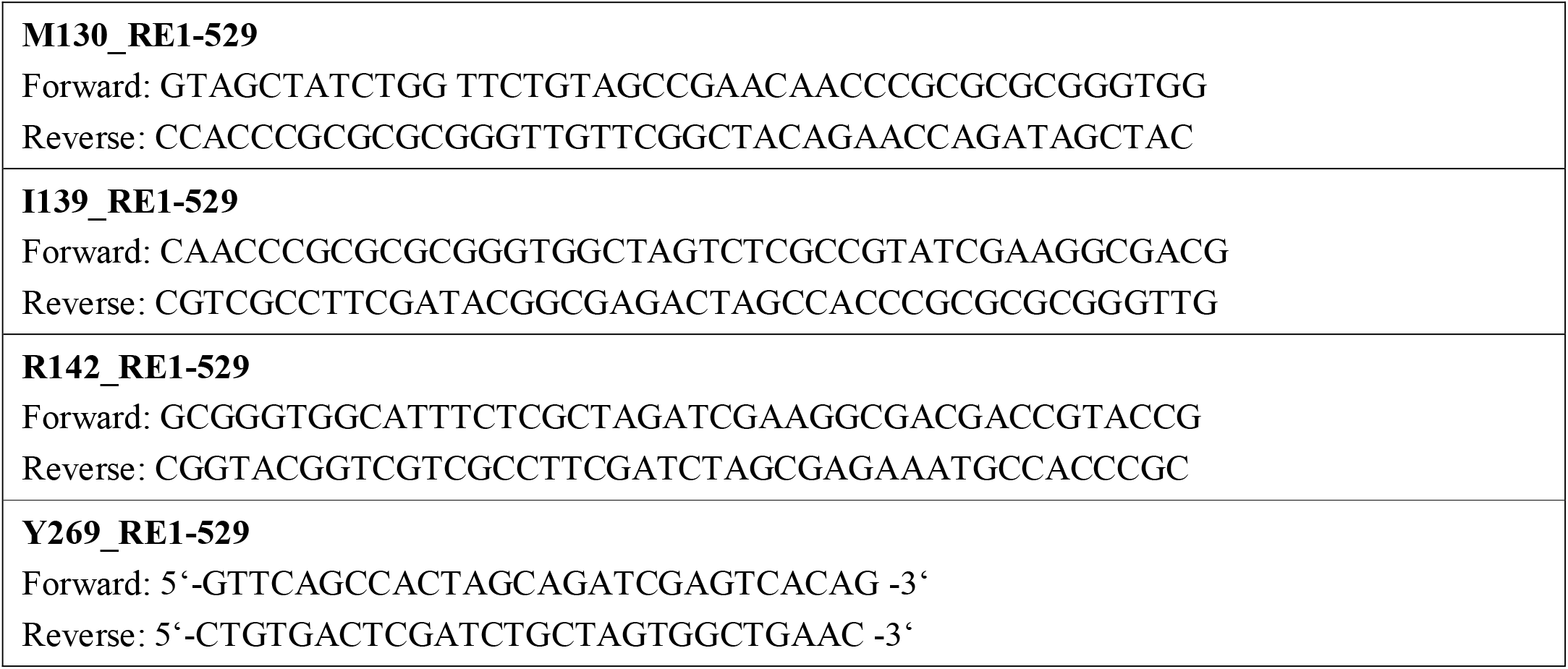
Primers for inserting conditional stop codons in RNase E

The p-azido phenylalanine incorporation was confirmed by biotinylation of azido group using EZ-link Phosphine-PEG3-Biotin (Thermo Fisher) (Agard et al. 2006; Saxon and Bertozzi 2000). Briefly, 50 µM of azido phenylalanine derivatives of RNase E NTD was reacted with 1 mM EZ-link Phosphine-PEG3-Biotin (x20 excess) at room temperature for 20 hrs. This allowed the phosphine group of EZ-link Phosphine-PEG3-Biotin to react with the azido group of p-azido phenylalanine, producing an aza-ylide intermediate (the Staudinger Reaction) (Saxon and Bertozzi 2000). Unbound biotin was removed by buffer exchanged into phosphate buffered saline by using Micro BioSpin-6 column concentrator, followed by concentrating to 50 µL. Samples were then loaded on SDS-PAGE gel and p-azido phenyl alanine was detected against anti-Biotin. Similar experiment was carried out with addition of reducing agent in PBS, resulting in a less intense band. While p-azido phenyl alanine derivatives showed band corresponding to NTD, the wild type NTD control did not show any band with the same procedure.

Cultures of transformed cells were grown in LB medium supplemented with carbenicillin (100 µg/mL), spectinomycin (40 µg/mL), arabinose (0.05% w/v), and p-AzidoPhe (1 mM) at 37 °C and 200 rpm. Cultures were induced by IPTG and cells were harvested by following the same procedure as used for NTD. P-AzidoPhe derivatives of NTD were purified by following the same procedure as used for NTD. The IMAC binding buffer was composed of 50 mM Phosphate buffer pH 7.9, 500 mM NaCl, 5 mM imidazole, 1 mM MgCl_2_, with elution buffer containing 500 mM imidazole. The size-exclusion buffer was composed of 50 mM Phosphate buffer pH 7.0, 500 mM NaCl, 10 mM MgCl_2_, 0.5 mM TCEP, 0.5 mM EDTA, 5% glycerol.

### Expression and purification of truncated degradosome

*E. coli* strain ENS134-10 was used to express RNase E 1-850 and full-length RhlB genes from the expression vector pRSF-DUET and full-length enolase from pET21b. Bacterial cultures, supplemented with 15 µg/mL kanamycin and 25 µg/mL carbenicillin were grown at 37° C until the OD_600_ reached 0.3-0.4 when protein production was induced by adding 1 mM IPTG. After overnight growth at 18° C, cells were harvested by centrifugation at 4200 g, 4° C for 30 minutes. Cells were resuspended in nickel-column buffer A (50 mM Tris pH 7.5, 1 M NaCl, 100 mM KCl, 5 mM imidazole, 10 mM MgCl_2_, 0.02% n-dodecyl β-D-maltoside (β-DDM)) and stored at -80° C until further use. Once thawed, the cells were supplemented with cOmplete EDTA-free protease inhibitor tablet (Roche), 1% Triton X-100, 1 mM TCEP, 1 mM PMSF, and 100 units of DNase I. Cells were lysed by passing the suspension through an EmulsiFlex-05 cell disruptor (Avestin) for 2-3 times at 10-15 kbar pressure. The lysate was clarified by centrifugation at 35000 g for 30 minutes and the supernatant was loaded onto a pre-equilibrated HiTrap Chelating HP column charged with nickel ions (GE Healthcare). The column was washed extensively with wash buffer (50 mM Tris pH 7.5, 1 M NaCl, 100 mM KCl, 100 mM imidazole, 10 mM MgCl_2_, 0.02% β-DDM), followed by elution of truncated degradosome by a gradient of elution buffer (50 mM Tris pH 7.5, 1 M NaCl, 100 mM KCl, 500 mM imidazole, 10 mM MgCl_2_, 0.02% β-DDM). Enriched fractions evaluated by SDS-PAGE were pooled together and passed through a cation exchange column (SP HP, GE Healthcare) which previously was equilibrated in a low-salt buffer (50 mM Tris pH 7.5, 50 mM NaCl, 10 mM KCl, 0.02% β-DDM). A linear gradient (0-50%) with ahigh-salt buffer (50 mM Tris pH 7.5, 2 M NaCl, 10 mM KCl, 0.02% β-DDM) was used to elute truncated degradosome. Desired fractions were pooled together, concentrated using 100 kDa MWCO concentrator and loaded onto a Superose6 10/300 size-exclusion column (GE Healthcare) equilibrated previously in storage buffer (50 mM HEPES pH 7.5, 400 mM NaCl, 100 mM KCl, 5 mM DTT, 0.02% β-DDM). Fractions containing the degradosome complex were flash frozen in liquid nitrogen and stored at -80 °C until further use.

### Expression and purification of full degradosome

*Escherichia coli* strain NCM3416 with a chromosomally strep-tagged RNase E was used to express the endogenous full-length RNA degradosome. Bacterial cultures were grown at 37° C in 2xYT media supplemented with 50 µg/mL kanamycin until the OD_600_ reached to 2.0 when protein production was induced by adding 1 mM IPTG. After overnight growth at 18° C, cells were harvested by centrifugation at 5020 g, 4° C for 30 minutes. Cells were resuspended in strep buffer A (50 mM Tris pH 7.5, 1 M NaCl, 100 mM KCl, 10 mM MgCl_2_, 0.02% β-DDM) and stored at -80° C until further use. Once thawed, the cells were supplemented with cOmplete EDTA-free protease inhibitor table (Roche), 1% Triton X-100, 1 mM TCEP, 1 mM PMSF, 100 units of DNase I, and 1 mg/mL lysozyme (Sigma). Cells were lysed by passing the suspension through an EmulsiFlex-05 cell disruptor (Avestin) for 2-3 times at 10-15 kbar pressure. The lysate was clarified by centrifugation at 35000 g for 30 minutes and the supernatant was passed through a 0.45 µ membrane filter before loading onto a pre-equilibrated HiTrapHP Strep column (GE Healthcare). The column was washed extensively with strep. Buffer A before the endogenous RNA degradosome was step-eluted with a strep buffer B (50 mM Tris pH 7.5, 200 mM NaCl, 100 mM KCl, 10 mM MgCl_2_, 0.02% β-DDM), followed by elution of truncated degradosome by elution buffer (50 mM Tris pH 7.5, 1 M NaCl, 100 mM KCl, 500 mM imidazole, 10 mM MgCl_2_, 0.02% β-DDM, 2.5 mM desbiotin(Sigma)). The best fractions were pooled and applied to a cation exchange column (HiTrap Heparin HP, GE Healthcare) equilibrated in a low-salt buffer (50 mM Tris pH 7.5, 50mM NaCl, 10 mM KCl, 0.02% β-DDM). A linear gradient (0-50%) with high-salt buffer (50 mM Tris pH 7.5, 2 M NaCl, 10 mM KCl, 0.02% β-DDM) was used to elute the full degradosome. Based on the purity of the eluted fractions, desired fractions were pooled together, concentrated down using 100 kDa MWCO concentrator and loaded onto a Superose6 10/300 size-exclusion column (GE Healthcare) equilibrated previously in storage buffer (50 mM HEPES pH 7.5, 400mM NaCl, 100 mM KCl, 5 mM DTT, 0.02% β-DDM). Desired fractions were flash frozen in liquid nitrogen and stored at -80 °C until further use.

### RNA preparation by *in vitro* transcription

RNAs were prepared by *in vitro* transcription. Plasmids with the 9S, RprA and GlmZ RNA genes were generously provided by A.J. Carpousis (CNRS, Toulouse), Kai Papenfort (Jena) and Boris Gorke (Vienna), respectively. First, genes were amplified by PCR using primers which were also adding T7 promoter. Next, RNA was synthesized from the PCR amplified product using T7 RNA polymerase at 37° C, followed by treating the reaction mixture with TURBO DNase for 15-20 minutes at 37° C. Finally, the RNA was purified by urea-PAGE followed by electroelution at 4° C and 100V (EluTrap, Whatman)(Bandyra, Wandzik, and Luisi 2018). In order to generate 5’-monophosphorylated RNA, rGMP was used in addition to rGTP (5:1 molar ratio) while keeping other reaction component and purification steps same as before(Bandyra, Wandzik, and Luisi 2018). For all RNAs, purity was checked in 8% urea-PAGE gel stained with SYBRgold RNA dye (Thermo Fisher).

### RNA degradation assays with pseudouridine substrates

polyA 20-mer, A20U and A20Ψ were obtained from Dharmacon. Oligoribonucleotides were 5□ labelled with ^32^ P using polynucleotide kinase (Fermentas), according to manufacturer instructions. Assays were carried out in reaction buffer (25 mM Tris-HCl pH 7.5, 50 mM NaCl, 50 mM KCl, 10 mM MgCl_2_, 1 mM DTT, 0.5 U/µL RNase OUT) at 37°C. 100 nM purified RNase E NTD was used for the reactions. Time course reactions were stopped at indicated time points by addition of STOP solution (20 mM EDTA, 2% w/v SDS). RNA loading dye (Thermo Fisher) was added to samples which were denatured (98°C, 2 min) and loaded onto polyacrylamide gels containing 7.5 M urea. Gels were dried and exposed to phosphor screens (GE Healthcare) and the signal analysed with TyphoonT 9400 (GE Healthcare).

### Kinetic assay

Ribonuclease cleavage of RNAs by RNase E was carried out at 30° C in the reaction buffer as above(Bandyra, Wandzik, and Luisi 2018). In case of time-course assay, samples were quenched at a predetermined time points by adding proteinase K mix (proteinase K in proteinase K buffer of 100 mM Tris-HCl pH 7.5, 150 mM NaCl, 12.5 mM EDTA, 1% SDS), followed by incubation at 50° C for 30 minutes. In the case of kinetic assay, RNA degradation was monitored against 10, 15, 20, 25, 50, 100, 125, 150, 200, 250, 300, 350, 400, 500, 600, 700 nM of the RNA while reaction was quenched as before by proteinase K within the linear range of the time-course curve. RNA samples were thereafter mixed with loading dye (Thermo Fisher), heated at 95° C for 2 minutes and loaded onto 8% urea-PAGE gel. The degradation products were visualized under UV transilluminator (GeneSnap, Syngene). To quantify, intensity of the reaction products was calculated using GeneTools (Syngene) with respect to a known amount of reference sample. Each kinetic parameter represents an average of three individual experiment.

## ACKNOWLEDGEMENTS

The work was supported by a Wellcome Trust Investigator award to BFL (200873/Z/16/Z). We thank Tom Dendooven, Giulia Paris and Steven Hardwick for helpful suggestions and advice. We thank A.J. Carpousis (CNRS, Toulouse) for expression plasmids and discussions.

## References

Agard, Nicholas J, Jeremy M Baskin, Jennifer A Prescher, Anderson Lo, and Carolyn R Bertozzi. 2006. “A Comparative Study of Bioorthogonal Reactions with Azides.” ACS Chemical Biology 1 (10): 644–48. https://doi.org/10.1021/cb6003228.

AÏt-Bara, Soraya, and Agamemnon J Carpousis. 2015. “RNA Degradosomes in Bacteria and Chloroplasts: Classification, Distribution and Evolution of RNase E Homologs.” Molecular Microbiology 97 (6): 1021–1135. https://doi.org/https://doi.org/10.1111/mmi.13095.

AÏt-Bara, Soraya, Agamemnon J Carpousis, and Yves Quentin. 2015. “RNase E in the γ-Proteobacteria: Conservation of Intrinsically Disordered Noncatalytic Region and Molecular Evolution of Microdomains.” Molecular Genetics and Genomics 290 (3): 847–62. https://doi.org/10.1007/s00438-014-0959-5.

Al-Husini, Nadra, Dylan T Tomares, Obaidah Bitar, W Seth Childers, and Jared M Schrader. 2018. “α-Proteobacterial RNA Degradosomes Assemble Liquid-Liquid Phase-Separated RNP Bodies.” Molecular Cell 71 (6): 1027-1039.e14. https://doi.org/https://doi.org/10.1016/j.molcel.2018.08.003.

Al-Husini, Nadra, Dylan T Tomares, Zechariah J Pfaffenberger, Nisansala S Muthunayake, Mohammad A Samad, Tiancheng Zuo, Obaidah Bitar, et al. 2020. “BR-Bodies Provide Selectively Permeable Condensates That Stimulate MRNA Decay and Prevent Release of Decay Intermediates.” Molecular Cell 78 (4): 670-682.e8. https://doi.org/https://doi.org/10.1016/j.molcel.2020.04.001.

Baker, Kristian E, and George A Mackie. 2003. “Ectopic RNase E Sites Promote Bypass of 5′-End-Dependent MRNA Decay in Escherichia Coli.” Molecular Microbiology 47 (1): 75–88. https://doi.org/https://doi.org/10.1046/j.1365-2958.2003.03292.x.

Bandyra, Katarzyna J, Marie Bouvier, Agamemnon J Carpousis, and Ben F Luisi. 2013. “The Social Fabric of the RNA Degradosome.” Biochimica et Biophysica Acta (BBA) - Gene Regulatory Mechanisms 1829 (6): 514–22. https://doi.org/https://doi.org/10.1016/j.bbagrm.2013.02.011.

Bandyra, Katarzyna J, Joanna M Wandzik, and Ben F Luisi. 2018. “Substrate Recognition and Autoinhibition in the Central Ribonuclease RNase E.” Molecular Cell 72 (2): 275–285.e4. https://doi.org/https://doi.org/10.1016/j.molcel.2018.08.039.

Boeynaems, Steven, Simon Alberti, Nicolas L Fawzi, Tanja Mittag, Magdalini Polymenidou, Frederic Rousseau, Joost Schymkowitz, et al. 2018. “Protein Phase Separation: A New Phase in Cell Biology.” Trends in Cell Biology 28 (6): 420–35. https://doi.org/https://doi.org/10.1016/j.tcb.2018.02.004.

Bruce, Heather A, Dijun Du, Dijana Matak-Vinkovic, Katarzyna J Bandyra, R William Broadhurst, Esther Martin, Frank Sobott, Alexander V Shkumatov, and Ben F Luisi. 2018. “Analysis of the Natively Unstructured RNA/Protein-Recognition Core in the Escherichia Coli RNA Degradosome and Its Interactions with Regulatory RNA/Hfq Complexes.” Nucleic Acids Research 46 (1): 387–402. https://doi.org/10.1093/nar/gkx1083.

Callaghan, Anastasia J, Maria Jose Marcaida, Jonathan A Stead, Kenneth J McDowall, William G Scott, and Ben F Luisi. 2005. “Structure of Escherichia Coli RNase E Catalytic Domain and Implications for RNA Turnover.” Nature 437 (7062): 1187–91. https://doi.org/10.1038/nature04084.

Campo, Cristian Del, Alexander Bartholomäus, Ivan Fedyunin, and Zoya Ignatova. 2015. “Secondary Structure across the Bacterial Transcriptome Reveals Versatile Roles in MRNA Regulation and Function.” PLOS Genetics 11 (10): e1005613. https://doi.org/10.1371/journal.pgen.1005613.

Carlile, Thomas M, Maria F Rojas-Duran, Boris Zinshteyn, Hakyung Shin, Kristen M Bartoli, and Wendy V Gilbert. 2014. “Pseudouridine Profiling Reveals Regulated MRNA Pseudouridylation in Yeast and Human Cells.” Nature 515 (7525): 143–46. https://doi.org/10.1038/nature13802.

Chandran, Vidya, Leonora Poljak, Nathalie F Vanzo, Anne Leroy, Ricardo Núñez Miguel, Juan Fernandez-Recio, James Parkinson, Christopher Burns, Agamemnon J Carpousis, and Ben F Luisi. 2007. “Recognition and Cooperation Between the ATP-Dependent RNA Helicase RhlB and Ribonuclease RNase E.” Journal of Molecular Biology 367 (1): 113–32. https://doi.org/https://doi.org/10.1016/j.jmb.2006.12.014.

Chao, Yanjie, Lei Li, Dylan Girodat, Konrad U Förstner, Nelly Said, Colin Corcoran, Michał Śmiga, et al. 2017. “In Vivo Cleavage Map Illuminates the Central Role of RNase E in Coding and Non-Coding RNA Pathways.” Molecular Cell 65 (1): 39–51. https://doi.org/https://doi.org/10.1016/j.molcel.2016.11.002.

Charette, Michael, and Michael W Gray. 2000. “Pseudouridine in RNA: What, Where, How, and Why.” IUBMB Life 49 (5): 341–51. https://doi.org/https://doi.org/10.1080/152165400410182.

Chatterjee, Debashree, Richard B Cooley, Chelsea D Boyd, Ryan A Mehl, George A O’Toole, and Holger Sondermann. 2014. “Mechanistic Insight into the Conserved Allosteric Regulation of Periplasmic Proteolysis by the Signaling Molecule Cyclic-Di-GMP.” Edited by Jon Clardy. ELife 3: e03650. https://doi.org/10.7554/eLife.03650.

Christiansen, Jan. 1988. “The 9S RNA Precursor of Escherichia Coli 5S RNA Has Three Structural Domains: Implications for Processing.” Nucleic Acids Research 16 (15): 7457–75. https://doi.org/10.1093/nar/16.15.7457.

Clarke, Justin E, Louise Kime, David Romero A., and Kenneth J McDowall. 2014. “Direct Entry by RNase E Is a Major Pathway for the Degradation and Processing of RNA in Escherichia Coli.” Nucleic Acids Research 42 (18): 11733–51. https://doi.org/10.1093/nar/gku808.

Cormack, Robert S, and George A Mackie. 1992. “Structural Requirements for the Processing of Escherichia Coli 5 S Ribosomal RNA by RNase E in Vitro.” Journal of Molecular Biology 228 (4): 1078–90. https://doi.org/https://doi.org/10.1016/0022-2836(92)90316-C.

Deana, Atilio, and Joel G Belasco. 2005. “Lost in Translation: The Influence of Ribosomes on Bacterial MRNA Decay.” Genes & Development 19 (21): 2526–33. https://doi.org/10.1101/gad.1348805.

Dreyfus, Marc B T -Progress in Molecular Biology and Translational Science. 2009. “Chapter 11 Killer and Protective Ribosomes.” In Molecular Biology of RNA Processing and Decay in Prokaryotes, 85:423–66. Academic Press. https://doi.org/https://doi.org/10.1016/S0079-6603(08)00811-8.

Durica-Mitic, Svetlana, and Boris Görke. 2019. “Feedback Regulation of Small RNA Processing by the Cleavage Product.” RNA Biology 16 (8): 1055–65. https://doi.org/10.1080/15476286.2019.1612693.

Garrey, Stephen M, Michaela Blech, Jenna L Riffell, Janet S Hankins, Leigh M Stickney, Melinda Diver, Ying-Han Roger Hsu, Vitharani Kunanithy, and George A Mackie. 2009. “Substrate Binding and Active Site Residues in RNases E and G.” Journal of Biological Chemistry 284 (46): 31843–50. https://doi.org/https://doi.org/10.1074/jbc.M109.063263.

Gonzalez, Grecia M, Svetlana Durica-Mitic, Steven W Hardwick, Martin C Moncrieffe, Marcus Resch, Piotr Neumann, Ralf Ficner, Boris Görke, and Ben F Luisi. 2017. “Structural Insights into RapZ-Mediated Regulation of Bacterial Amino-Sugar Metabolism.” Nucleic Acids Research 45 (18): 10845–60. https://doi.org/10.1093/nar/gkx732.

Göpel, Yvonne, Kai Papenfort, Birte Reichenbach, Jörg Vogel, and Boris Görke. 2013. “Targeted Decay of a Regulatory Small RNA by an Adaptor Protein for RNase E and Counteraction by an Anti-Adaptor RNA.” Genes & Development 27 (5): 552–64. https://doi.org/10.1101/gad.210112.112.

Guzikowski, Anna R, Yang S Chen, and Brian M Zid. 2019. “Stress-Induced MRNP Granules: Form and Function of Processing Bodies and Stress Granules.” WIREs RNA 10 (3): e1524. https://doi.org/https://doi.org/10.1002/wrna.1524.

Guzzi, Nicola, Maciej CieŚla, Phuong Cao Thi Ngoc, Stefan Lang, Sonali Arora, Marios Dimitriou, Kristyna Pimková, et al. 2018. “Pseudouridylation of TRNA-Derived Fragments Steers Translational Control in Stem Cells.” Cell 173 (5): 1204–1216.e26. https://doi.org/https://doi.org/10.1016/j.cell.2018.03.008.

Hadjeras, Lydia, Leonora Poljak, Marie Bouvier, Quentin Morin-Ogier, Isabelle Canal, Muriel Cocaign-Bousquet, Laurence Girbal, and Agamemnon J Carpousis. 2019. “Detachment of the RNA Degradosome from the Inner Membrane of Escherichia Coli Results in a Global Slowdown of MRNA Degradation, Proteolysis of RNase E and Increased Turnover of Ribosome-Free Transcripts.” Molecular Microbiology 111 (6): 1715–31. https://doi.org/https://doi.org/10.1111/mmi.14248.

Kalamorz, Falk, Birte Reichenbach, Walter März, Bodo Rak, and Boris Görke. 2007. “Feedback Control of Glucosamine-6-Phosphate Synthase GlmS Expression Depends on the Small RNA GlmZ and Involves the Novel Protein YhbJ in Escherichia Coli.” Molecular Microbiology 65 (6): 1518–33. https://doi.org/https://doi.org/10.1111/j.1365-2958.2007.05888.x.

Khemici, Vanessa, and Agamemnon J Carpousis. 2004. “The RNA Degradosome and Poly(A) Polymerase of Escherichia Coli Are Required in Vivo for the Degradation of Small MRNA Decay Intermediates Containing REP-Stabilizers.” Molecular Microbiology 51 (3): 777–90. https://doi.org/https://doi.org/10.1046/j.1365-2958.2003.03862.x.

Kime, Louise, Justin E Clarke, David Romero A., Jane A Grasby, and Kenneth J McDowall. 2014. “Adjacent Single-Stranded Regions Mediate Processing of TRNA Precursors by RNase E Direct Entry.” Nucleic Acids Research 42 (7): 4577–89. https://doi.org/10.1093/nar/gkt1403.

Kime, Louise, Stefanie S Jourdan, Jonathan A Stead, Ana Hidalgo-Sastre, and Kenneth J McDowall. 2010. “Rapid Cleavage of RNA by RNase E in the Absence of 5′ Monophosphate Stimulation.” Molecular Microbiology 76 (3): 590–604. https://doi.org/https://doi.org/10.1111/j.1365-2958.2009.06935.x.

Koslover, Daniel J, Anastasia J Callaghan, Maria J Marcaida, Elspeth F Garman, Monika Martick, William G Scott, and Ben F Luisi. 2008. “The Crystal Structure of the Escherichia Coli RNase E Apoprotein and a Mechanism for RNA Degradation.” Structure 16 (8): 1238–44. https://doi.org/https://doi.org/10.1016/j.str.2008.04.017.

Leroy, Anne, Nathalie F Vanzo, Sandra Sousa, Marc Dreyfus, and Agamemnon J Carpousis. 2002. “Function in Escherichia Coli of the Non-Catalytic Part of RNase E: Role in the Degradation of Ribosome-Free MRNA.” Molecular Microbiology 45 (5): 1231–43. https://doi.org/https://doi.org/10.1046/j.1365-2958.2002.03104.x.

Lin, Yuan, David S.W. Protter, Michael K. Rosen, and Roy Parker. 2015. “Formation and Maturation of Phase-Separated Liquid Droplets by RNA-Binding Proteins.” Molecular Cell 60 (2): 208–19. https://doi.org/https://doi.org/10.1016/j.molcel.2015.08.018.

Lorenz, Ronny, Stephan H Bernhart, Christian Höner zu Siederdissen, Hakim Tafer, Christoph Flamm, Peter F Stadler, and Ivo L Hofacker. 2011. “ViennaRNA Package 2.0.” Algorithms for Molecular Biology 6 (1): 26. https://doi.org/10.1186/1748-7188-6-26.

Mackie, George A. 1998. “Ribonuclease E Is a 5′-End-Dependent Endonuclease.” Nature 395 (6703): 720–24. https://doi.org/10.1038/27246.

Mackie, George A.. 2013. “RNase E: At the Interface of Bacterial RNA Processing and Decay.” Nature Reviews Microbiology 11 (1): 45–57. https://doi.org/10.1038/nrmicro2930.

Marcaida, Maria Jose, Mark A DePristo, Vidya Chandran, Agamemnon J Carpousis, and Ben F Luisi. 2006. “The RNA Degradosome: Life in the Fast Lane of Adaptive Molecular Evolution.” Trends in Biochemical Sciences 31 (7): 359–65. https://doi.org/10.1016/j.tibs.2006.05.005.

Moffitt, Jeffrey R, Shristi Pandey, Alistair N Boettiger, Siyuan Wang, and Xiaowei Zhuang. 2016. “Spatial Organization Shapes the Turnover of a Bacterial Transcriptome.” Edited by Rachel Green. ELife 5: e13065. https://doi.org/10.7554/eLife.13065.

Nott, Timothy J., Evangelia Petsalaki, Patrick Farber, Dylan Jervis, Eden Fussner, Anne Plochowietz, Timothy D Craggs, et al. 2015. “Phase Transition of a Disordered Nuage Protein Generates Environmentally Responsive Membraneless Organelles.” Molecular Cell 57 (5): 936–47. https://doi.org/https://doi.org/10.1016/j.molcel.2015.01.013.

Pereira, Joana, and Andrei N Lupas. 2018. “The Ancestral KH Peptide at the Root of a Domain Family with Three Different Folds.” Bioinformatics 34 (23): 3961–65. https://doi.org/10.1093/bioinformatics/bty480.

Redko, Yulia, Mark R Tock, Chris J Adams, Vladimir R Kaberdin, Jane A Grasby, and Kenneth J McDowall. 2003. “Determination of the Catalytic Parameters of the N-Terminal Half of Escherichia Coli Ribonuclease E and the Identification of Critical Functional Groups in RNA Substrates*.” Journal of Biological Chemistry 278 (45): 44001–8. https://doi.org/https://doi.org/10.1074/jbc.M306760200.

Richards, Jamie, and Joel G Belasco. 2021. “Widespread Protection of RNA Cleavage Sites by a Riboswitch Aptamer That Folds as a Compact Obstacle to Scanning by RNase E.” Molecular Cell 81 (1): 127-138.e4. https://doi.org/https://doi.org/10.1016/j.molcel.2020.10.025.

Saxon, Eliana, and Carolyn R Bertozzi. 2000. “Cell Surface Engineering by a Modified Staudinger Reaction.” Science 287 (5460): 2007LP–2010. https://doi.org/10.1126/science.287.5460.2007.

Strahl, Henrik, Catherine Turlan, Syma Khalid, Peter J Bond, Jean-Marie Kebalo, Pascale Peyron, Leonora Poljak, et al. 2015. “Membrane Recognition and Dynamics of the RNA Degradosome.” PLOS Genetics 11 (2): e1004961. https://doi.org/10.1371/journal.pgen.1004961.

Sulthana, Shaheen, Georgeta N Basturea, and Murray P Deutscher. 2016. “Elucidation of Pathways of Ribosomal RNA Degradation: An Essential Role for RNase E.” RNA 22 (8): 1163–71. https://doi.org/10.1261/rna.056275.116.

Thompson, Katharine J, Jeff Zong, and George A Mackie. 2015. “Altering the Divalent Metal Ion Preference of RNase E.” Edited by R L Gourse. Journal of Bacteriology 197 (3): 477LP–482. https://doi.org/10.1128/JB.02372-14.

Tsai, Yi-Chun, Dijun Du, Lilianha Domínguez-Malfavón, Daniela Dimastrogiovanni, Jonathan Cross, Anastasia J Callaghan, Jaime García-Mena, and Ben F Luisi. 2012. “Recognition of the 70S Ribosome and Polysome by the RNA Degradosome in Escherichia Coli.” Nucleic Acids Research 40 (20): 10417–31. https://doi.org/10.1093/nar/gks739.

Updegrove, Taylor B, Andrew B Kouse, Katarzyna J Bandyra, and Gisela Storz. 2019. “Stem-Loops Direct Precise Processing of 3′ UTR-Derived Small RNA MicL.” Nucleic Acids Research 47 (3): 1482–92. https://doi.org/10.1093/nar/gky1175.

Urban, Johannes H, and Jörg Vogel. 2008. “Two Seemingly Homologous Noncoding RNAs Act Hierarchically to Activate GlmS MRNA Translation.” PLOS Biology 6 (3): e64. https://doi.org/10.1371/journal.pbio.0060064.

